# Perinatal exposures and upper respiratory tract microbiome composition are associated with age at first acute otitis media episode

**DOI:** 10.64898/2026.07.13.738158

**Authors:** Corbin R. Hite, Congwen Zhao, Kate Hoffman, Jillian H. Hurst

## Abstract

Acute otitis media (AOM) is the most common bacterial infection of childhood and the leading indication for antibiotic prescriptions and healthcare consultations globally. Colonization of the upper respiratory tract (URT) microbiome by bacterial respiratory pathogens precedes AOM episodes; however, the factors that influence colonization susceptibility and subsequent AOM are not well understood. We hypothesized that perinatal exposures, including mode of delivery, intrapartum antibiotic exposure, and infant feeding influence the composition of the URT microbiome at birth, modifying risk of AOM in infancy. We characterized the URT microbiome in nasopharyngeal swabs collected from 163 infants at birth. Swabs were generally collected within two days of delivery (median [IQR] collection time: 25 [17, 45] hours) and microbiome composition was evaluated with 16S rRNA V4 sequencing. Exposures evaluated included birth mode, intrapartum antibiotic exposures, and feeding type at hospital discharge. AOM episodes were identified through electronic health records data. We built Cox proportional hazards models to determine if perinatal exposures and/or microbiome characteristics at birth were associated with the time to first AOM episode in the first two years of life. URT microbiome diversity and composition were associated with feeding type at hospital discharge, wherein exclusive formula feeding was associated with increased diversity and the presence of *Staphylococcus* and *Haemophilus* spp. Increased URT microbiome diversity was associated with younger age at first AOM episode. Our findings suggest that perinatal exposures may influence the composition of the birth URT microbiome, and that this early composition may be related to AOM susceptibility in infancy.

**Importance:** Ear infections are the most common bacterial infection of childhood and the leading indication for healthcare consultation and antibiotic receipt. Previous studies have demonstrated that the microbes that inhabit the upper respiratory tract, known as the microbiome, influence risk of ear infection. This study sought to understand how exposures around the time of birth, including delivery type, maternal antibiotic exposures, and infant feeding, influence the development of the infant microbiome, and in turn, how the microbiome is related to ear infections. An analysis of nasal swabs collected from infants shortly after birth demonstrated that increased microbial diversity is associated with earlier age at first ear infection episode. Overall, this study demonstrates that exposures in early life influence respiratory microbiome development, which contributes to infection susceptibility.

## Introduction

Acute otitis media (AOM) is the most common bacterial infection in children globally, with over 700 million cases each year (1). Moreover, AOM is the leading indication for healthcare consultation and antibiotic prescriptions (2, 3). Previous work has linked perinatal exposures, occurring before or during delivery and in the several weeks following, with lasting impacts on AOM risk in childhood. Several studies have suggested that birth via cesarean section is associated with a slight increase in risk of AOM in early life (4–6). Maternal antibiotic use during pregnancy was also found to be associated with an increased risk of AOM (7–10). Administration of intrapartum antibiotics during delivery has been associated with increased risk for overall infection in infancy, but the relationship with AOM remains understudied (11). Breastfed infants have been shown to be at lower risk of AOM in early childhood, with one study finding that exclusive or supplemental formula-feeding in the first 6 months of life increases risk of AOM in early childhood (8, 12–16). Though these studies have provided some insights into potential risk factors for AOM, our understanding of these factors and the underlying mechanisms remain incomplete.

The perinatal and early life exposures described above have been linked to changes to the community of microbes that reside within the human upper respiratory tract (URT), referred to as the URT microbiome (17–19). The URT microbiome is a modifiable aspect of host physiology that plays a critical role in the development of the immune system and in mediating resistance to pathogen colonization. The composition of the URT microbiome undergoes rapid development during the first few years of life and continues to evolve through late adolescence (20, 21). In early life, URT microbiome composition is influenced by infant feeding practices, exposure to congregate care settings or crowding, antibiotic receipt, and tobacco smoke exposure (22, 23). Thus, early life exposures influence URT microbiome composition and likely influence its effects on early life infections, including AOM (24, 25).

Increasing evidence has underscored the role of the URT microbiome in AOM development. AOM episodes are preceded by bacterial pathogen colonization of the URT (26). Importantly, the composition of the URT microbiome can directly modulate colonization within this niche and subsequent AOM risk (27, 28). For example, Xu and colleagues found that children who were “otitis-prone” had decreased URT microbiome diversity compared to AOM-free children (29). Notably, lower diversity of the URT microbiome was associated with higher rates of colonization by otopathogens such as *S. pneumoniae*, *H. influenzae*, and *M. catarrhalis* (30, 31). In contrast, the presence of *Dolosigranulum* and *Corynebacterium* were found to reduce the risk of AOM development (32, 33). Taken together, these findings suggest that URT microbiome composition is associated with AOM risk and may partially explain interindividual differences in susceptibility among young children. Further, while evidence suggests associations with both perinatal exposures and URT microbiome composition with early life AOM, little is known about how perinatal exposures act on the development of the early life URT microbiome to potentially influence AOM susceptibility.

To gain a better understanding of the role of perinatal exposures and the early life URT microbiome in AOM, we evaluated birth URT microbiome composition in a cohort of 163 infants enrolled in a prospective birth cohort study. We specifically assessed associations between microbiome composition and perinatal exposures, including delivery type, exposure to intrapartum antibiotics, and infant feeding practices. Finally, we determined how these perinatal exposures and features of the URT microbiome at the time of birth influenced timing of the first AOM episode in the first two years of life.

## Materials and Methods

### Study population

The study was performed using samples and data from HOPE 1000, a prospective pregnancy cohort based at Duke University. The cohort includes 525 mother-infant pairs enrolled from 2018 to 2025. Eligible pregnant participants were 18 years of age or older, had a singleton pregnancy, received prenatal care through the Duke University Health System (DUHS), and intended to remain in the area for at least two years and have their infants receive pediatric care through DUHS. Mothers were recruited during pregnancy up to 24 weeks gestation and had study visits at each enrolled trimester, delivery, and 6-8 weeks postpartum. Infants were automatically enrolled at birth and had study visits at delivery and over the first two years of life. During each study visit, participants had biospecimens collected and completed demographic and behavioral surveys. All participants were consented to passive follow-up via electronic health records (EHR) data until the child was 5 years of age. Samples used for the current study were collected from 2020 to 2023.

### Specimen collection and processing

A nasopharyngeal swab was collected from infants at delivery using nylon flocked swabs (Copan Italia, Brescia, Italy) stored in RNAProtect (Qiagen, Hilden, Germany). Swabs were collected before the infant was discharged from the hospital. The age in hours at time of swab collection was calculated. Swab samples were aliquoted and stored at −80 °C before being thawed for DNA extraction and sequencing.

### 16S rRNA gene amplicon sequencing and analysis

The Duke Microbiome Core Facility extracted DNA from nasal swab samples using a modified Macherey-Nagel NucleoSpin Tissue XS Kit (Takara Bio, San Jose, CA) with MN Bead Tubes Type B (Takara Bio, San Jose, CA). Bacterial community composition in isolated DNA samples was characterized by amplification of the V4 variable region of the 16S rRNA gene by polymerase chain reaction (PCR) using the forward primer 515 (5’-GTGCCAGCMGCCGCGGTAA-3’) and reverse primer 806 (5’-GGACTACHVGGGTWTCTAAT-3’) from the Earth Microbiome Project protocol (34). The reverse primers (806R) carry unique barcodes that allow for multiplexed sequencing. PCR was performed using Phusion Plus PCR Master Mix (ThermoFisher Scientific, Waltham, MA) for a total of 35 cycles. PCR products were purified with AMPure XP Beads (Beckman Coulter, Brea, CA), and concentrations were assessed using a Qubit dsDNA HS assay kit (ThermoFisher Scientific, Waltham, MA) and a Promega GloMax plate reader (Promega Corporation, Madison, WI). Equimolar 16S rRNA PCR products from all samples were pooled prior to sequencing. Sequencing was performed by the Duke Sequencing and Genomic Technologies shared resource on an Illumina MiSeq instrument configured for 250 base-pair paired-end sequencing runs (Illumina, San Diego, CA). Negative controls were included in all stages of extraction and PCR. Sequencing reads were processed through the *DADA2* package (35) and amplicon sequence variant (ASV) taxonomy was assigned based on the expanded Human Oral Microbiome Database (eHOMD) version 15.1 (36). Due to the low biomass of infant nasopharyngeal swabs, we anticipated that our samples would be especially influenced by contamination. A list of reagent contaminants was compiled from ASVs detected in any of the negative controls and those identified via the *decontam* package (Frequency method; Threshold = 0.1)(37). Contaminant ASVs (n = 554) were removed from all samples in the study (**Table S1)**. Samples with a read-depth of less than 1,000 reads after contaminant removal were excluded from further analysis. These quality measures resulted in 163 birth URT microbiome samples with a median (interquartile range [IQR]) of 6,004 (2,842–13,482) reads. Reads were mapped to 1,734 total ASVs representing 136 genera from 11 phyla.

### EHR data abstraction

We used the Duke Clinical Research Datamart to abstract participant EHR data (38). Data included types and dates of encounters, infant age at encounter, maternal medications administered during delivery, mode of delivery, ICD-10 diagnosis codes, NICU admission, and well-child visit attendance. We defined infant intrapartum antibiotic exposure as maternal antibiotic receipt during the delivery encounter but prior to the time of birth. Breastfeeding status was indicated as the feeding type at time of infant hospital discharge after delivery. We recorded the presence and timing of AOM episodes in the infant’s first 2 years of life based on ICD-10 codes (**Table S2**).

To further assess the influence of early feeding practices on AOM outcomes, we abstracted EHR data for a large retrospective cohort of children who received regular well child care through DUHS over the first two years of life (n = 4,378). Children were eligible for inclusion if they were born at a DUHS hospital between January 1, 2021, and June 1, 2023, and had attended 6 or more well child visits by age 2. We additionally limited the cohort to infants with at least four infant feeding records, as documented through the Milk Box flow sheet, which is used by DUHS pediatric primary care providers to capture infant feeding practices from birth to age 2 (39). For each infant included in the cohort, we abstracted the number of months of exclusive breastfeeding, mixed breast and formula feeding, and exclusive formula feeding. We also abstracted the time of first AOM episode (if any), the number of well child visits attended, mode of delivery, intrapartum antibiotic exposure, and NICU admission. We excluded children with chronic conditions that could influence their susceptibility to AOM, including children with genetic disorders (e.g., trisomy 21), cleft lip and/or palate, or immune deficiencies (**Table S2**).

### Statistical analyses

We used Chi-squared or Fisher’s exact tests (categorical variables) and Wilcoxon rank-sum or Kruskal–Wallis tests (continuous variables) to evaluate differences in sociodemographic and clinical characteristics between modes of delivery, intrapartum antibiotic exposures, and feeding practices. We evaluated microbiome alpha diversity and richness using the *phyloseq* package (40). We described the relative abundance of the top 30 taxa identified across the cohort as a percentage of each taxon’s reads over total sample reads. The mean relative abundance was computed as the mean proportion for each taxa across the group to be summarized and the total abundances were re-normalized to a proportion out of 1.0. We quantified the overall microbiome composition via Bray-Curtis dissimilarity, which was computed using a principal component ordination in the *vegan* package. We compared Bray-Curtis dissimilarity between exposure groups with PERMANOVA using the adonis2 function in the *vegan* package (41).

A presence/absence approach was used to determine if the proportion of samples in which specific taxa were detected varied between groups. The taxa of interest were chosen due to their high prevalence in the URT microbiome and previous literature suggesting their involvement in respiratory health (29, 32, 33). These genera include *Streptococcus, Staphylococcus, Haemophilus, Dolosigranulum, Moraxella, and Corynebacterium.* We used Fisher’s exact tests to evaluate for differences in the proportions across groups.

We built multivariate Cox proportional hazards models (42) to evaluate associations between infant exposures, microbiome characteristics, and the time to first AOM episode in the first two years of life. We adjusted all models for race/ethnicity, NICU admission, and well-child visit attendance. To determine the impacts of infant feeding practices on AOM susceptibility in early life, we used data from a large retrospective EHR cohort (described above). A Cox proportional hazards model was used to evaluate for associations between infant feeding practices and time to first AOM episode, adjusting for race/ethnicity, gestational age at delivery, intrapartum antibiotic exposure, mode of delivery, NICU admission, and well-child visit attendance. All analyses were conducted using R version 4.4.2 (45)

We conducted an unsupervised clustering analysis using the Calinski-Harabasz index and k-medoids clustering to classify samples into distinct microbiome profiles (43, 44). We used Chi-squared or Fisher’s exact tests (categorical variables) and Wilcoxon rank-sum or Kruskal–Wallis tests (continuous variables) to evaluate for associations between infant characteristics and exposures with microbiome profiles. The significance threshold for all statistical tests was set at α = 0.05.

## Results

### Study population

We included 163 infants with birth microbiome samples that passed quality controls. Slightly under half (47%) were female, and the median gestational age at birth was 39 (interquartile range [IQR], 37.3, 39.4) weeks. A little less than half (43%) of the cohort was born via Cesarean section, 29% were exposed to intrapartum antibiotics, and 5% were exclusively formula fed at the time of hospital discharge. Children were a median of 25 (IQR: 17, 45) hours old at time of nasopharyngeal swab collection (**Table 1**). Children born via Cesarean section were significantly older at time of nasopharyngeal swab collection compared to those born vaginally (vaginal, median (IQR): 22 (12, 31) hours; Cesarean section, median (IQR): 39 (22, 58) hours; Wilcoxon rank-sum; p < 0.001; **Table S3**). The cohort included 50 (31%) children born to mothers who tested positive for group-B *Streptococcus* (GBS) colonization, of which 34 (68%) were exposed to intrapartum antibiotics (**Table S4**).

**Table 1.** Characteristics of the study population.

| Table 1. Characteristics of the study population |  |  |
| --- | --- | --- |
| Infant characteristics (n=163) |  | n (%) or median (IQR) |
| Age at sample collection (hours), median (IQR) |  | 25 (17 – 45) |
| Female sex |  | 76 (47%) |
| Birth weight (g), median (IQR) |  | 3,320 (2,920-3,615) |
| Gestational age at birth (weeks), median (IQR) |  | 39.0 (37.3-39.4) |
| Race |  |  |
|  | Non-Hispanic White | 68 (42%) |
|  | Non-Hispanic Black | 56 (34%) |
|  | Non-Hispanic Asian | 9 (5.5%) |
|  | Hispanic | 7 (4.3%) |
|  | Other/Multiple Races | 23 (14%) |
| Annual household income |  |  |
| | Under \$50,000 | 48 (30%) |
| | \$50,000 - \$149,000 | 58 (36%) |
| | \$150,000+ | 41 (25%) |
|  | Not reported | 14 (8.7%) |
| Birth Year |  |  |
|  | 2020 or 2021 | 32 (20%) |
|  | 2022 | 81 (50%) |
|  | 2023 | 50 (31%) |
| Cesarean delivery |  | 70 (43%) |
| Maternal GBS positive status |  | 50 (30%) |
| Intrapartum antibiotic exposure |  | 48 (29%) |
| Formula feeding at discharge |  | 8 (5%) |
| NICU admission |  | 33 (20%) |
| GBS, Group-B *Streptococcus*<br>NICU, Neonatal intensive care unit |  |  |

### Associations between birth URT microbiome composition and perinatal exposures

We first assessed associations between URT microbiome composition and perinatal exposures, including mode of delivery, intrapartum antibiotic exposure, and infant feeding type at discharge (**Figure 1**). We did not identify any associations between mode of delivery or intrapartum antibiotic exposure with any measure of microbiome composition, including richness (vaginal, median (IQR): 21 (13, 33); Cesarean section, median (IQR): 21 (14, 33); **Figure 1A**; unexposed to antibiotics (IQR): 22 (14.74, 31.25); exposed to antibiotics (IQR): 18 (12, 36); **Figure 1C**), the Shannon diversity index (vaginal, median (IQR): 2.27 (1.70, 2.66); Cesarean section, median (IQR): 2.39 (1.74, 2.74); **Figure 1B**; unexposed to intrapartum antibiotics (IQR): 2.39 (1.73, 2.75); exposed to intrapartum antibiotics (IQR): 2.24 (1.73, 2.67); **Figure 1D**), presence of taxa of interest (**Figure 2A, 2C**) or Bray-Curtis dissimilarity (PERMANOVA; mode of delivery, F_pseudo_: 1.28; **Figure 2B**; intrapartum antibiotic exposure, F_pseudo_: 0.58; **Figure 1D**). There were significant differences in microbiome composition between infants who were breastfed versus formula fed at the time of hospital discharge. Formula-fed infants had a significant increase in the observed number of ASVs within the URT microbiome compared to infants who were breastfed (breastfed, median (IQR): 21 (13, 32.5) versus formula fed, median (IQR): 35.5 (30.25, 49), Wilcoxon rank sum, p = 0.009, **Figure 1E**). Similarly, we observed a significant increase in the Shannon diversity index of formula-fed infants compared to breastfed infants (breastfed, median (IQR) 2.29 (1.70, 2.66) versus formula fed, median (IQR): 2.90 (2.84, 3.07), Wilcoxon rank sum, p = 0.003, **Figure 1F**), and a significant increase in the proportion of samples in which taxa assigned to *Haemophilus* (breastfed proportion detected: 0.28, formula fed proportion detected: 0.75, Fisher’s exact test, p = 0.005, **Figure 2E**) and *Staphylococcus* (breastfed proportion detected: 0.28, formula fed proportion detected: 0.63, Fisher’s exact test, p = 0.040) were detected when compared to samples from breastfed infants. A PERMANOVA comparing Bray-Curtis dissimilarity did not identify differences in the overall microbiome composition by feeding type (F_pseudo_: 1.19; **Figure 2F**).

**Figure 1.**
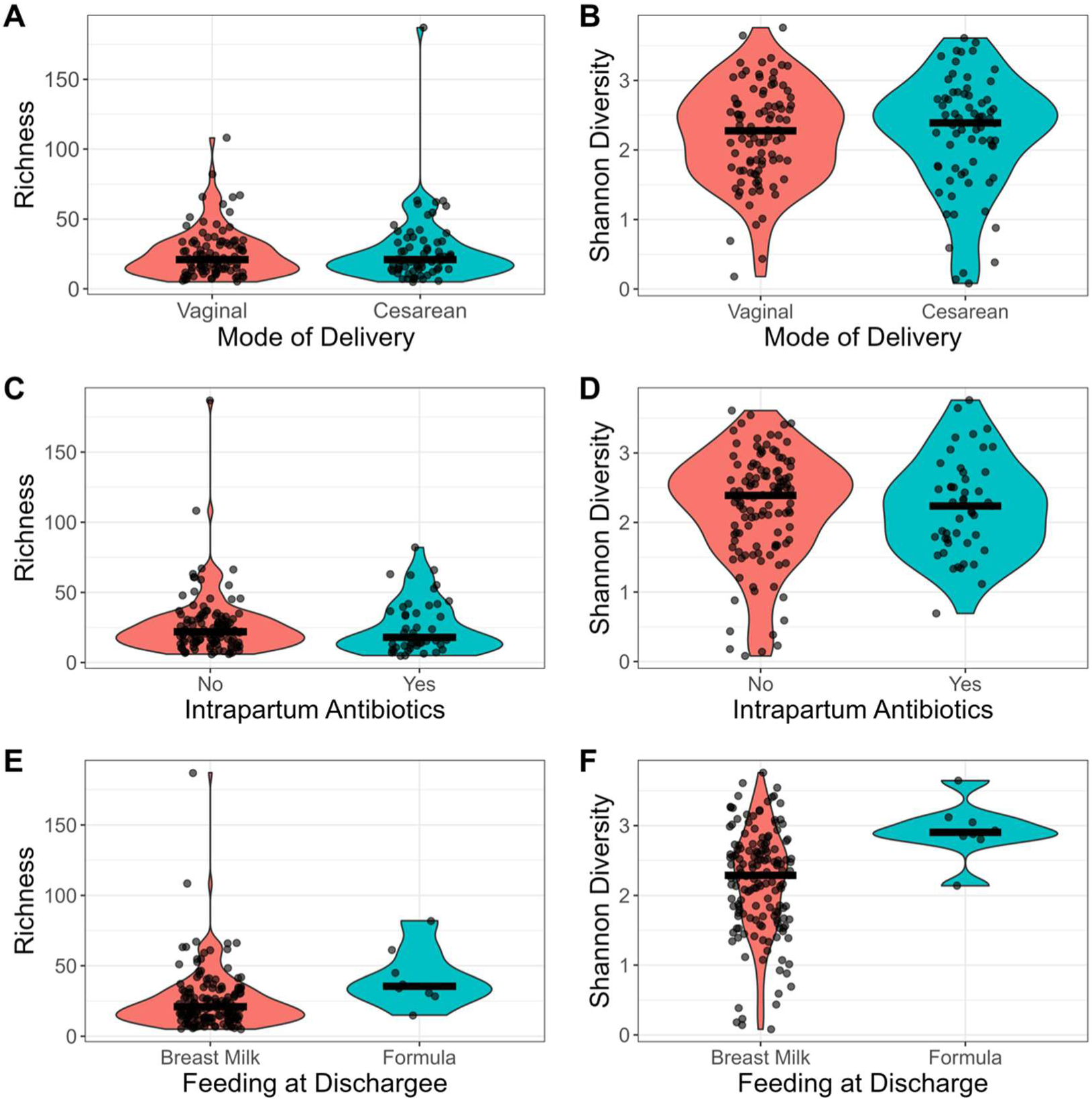
The influence of perinatal exposures on URT microbiome alpha diversity. We evaluated for associations in within-sample (alpha) diversity, including richness, or observed ASVs (**A, C, E**) and the Shannon diversity (**B, D, F**) of the URT microbiome. Samples were evaluated by mode of delivery (vaginal birth: n = 93, Cesarean section: n = 70) (**A, B**), intrapartum antibiotic exposure (unexposed n = 116, exposed n = 47) (**C, D**), and infant feeding type at hospital discharge (breastfed: n = 155, formula fed: n = 8) (**E, F**). Differences were evaluated by Wilcoxon rank sum test.

**Figure 2.**
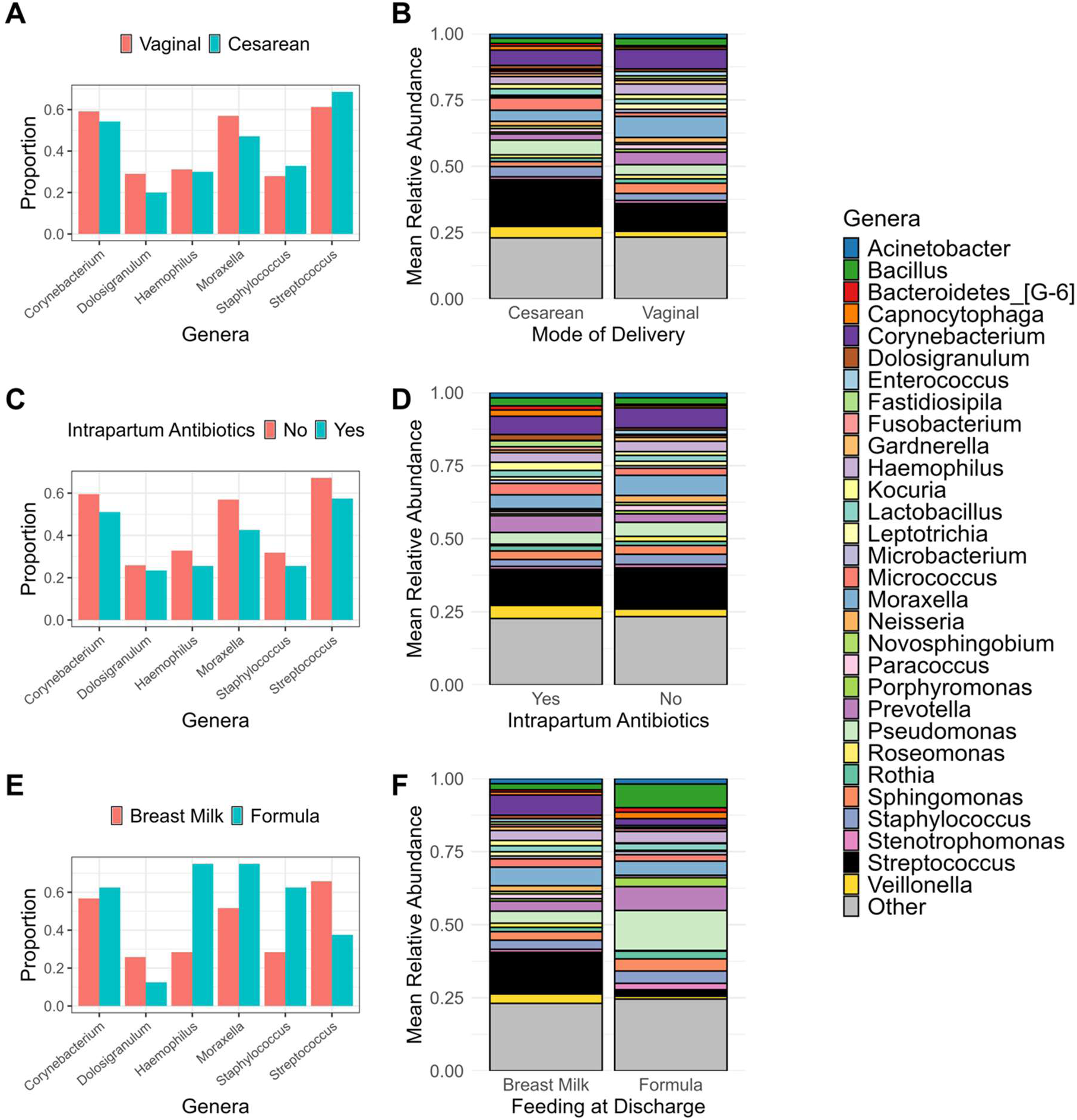
Associations between perinatal exposures and URT microbiome composition. We evaluated for differences in birth URT microbiome composition by (**A, B**) mode of delivery (vaginal birth: n = 93, Cesarean section: n = 70), (**C, D**) intrapartum antibiotic exposure (unexposed n = 116, exposed n = 47), and (**E, F**) feeding type at discharge (breastfed: n = 155, formula fed: n = 8). (**A, C, E**) Bar plots show the difference between the proportion of samples in which genera of interest were detected in the URT microbiome, with significant differences evaluated by Fisher’s exact tests. (**B, D, F**) Stacked bar plots display the relative abundance of the top 30 genera identified in the URT microbiome. Differences in the overall composition of the URT microbiome was assessed using PERMANOVA of Bray-Curtis dissimilarity by exposure group.

### Associations between perinatal exposures, URT microbiome features, and time to first AOM episode

We next assessed associations between perinatal exposures and time to first AOM episode in the first two years of life. We built Cox proportional hazards models using mode of delivery (**Figure 3A**), intrapartum antibiotic exposures (**Figure 3B**), and feeding type at hospital discharge (**Figure 3C**) as the exposures of interest. All models were adjusted for income, NICU admission, and well child visit attendance as a measure of observability. We did not identify any associations between time to first AOM episode and mode of delivery (Cesarean section, HR [CI], 1.44 [0.89, 2.32]) or intrapartum antibiotic administration (Exposed, HR [CI], 0.91 [0.53, 1.57]) within the HOPE cohort (**Figure 2A, 2B**). We observed a slight but non-significant decrease in HR for infants who were formula fed (HR [CI], 0.78 [0.24, 2.57]; **Figure 2C, 2D**); however, only 8 infants in our cohort were formula fed at the time of hospital discharge, indicating a lack of power to detect potential differences. Given this small sample size, we decided to further explore the relationship between infant feeding type and time to first AOM episode using a large, retrospective electronic health records (EHR) dataset. We constructed a retrospective cohort of infants with data from birth to age 2 using EHR data from Duke University Health System. We identified 4,378 infants born in the DUHS who met the criteria for inclusion in an analysis of feeding type and AOM. Slightly under half of infants in the cohort were female (49%) and the median gestational age at birth was 39 (interquartile range [IQR], 37.9-39.9) weeks (**Table S4**). Around a third of infants (33%) were born via cesarean section and 10% were exposed to intrapartum antibiotics. We assigned feeding status as a dichotomous variable indicating if an infant received any breastmilk in the first six months of life or was exclusively formula fed. Slightly over 18% of the cohort was exclusively formula-fed in the first 6 months of life. Nearly half of infants (46.8%) had a documented AOM episode by age 2. We built a Cox proportional hazards model to evaluate associations between exclusive formula feeding and AOM in the first two years of life, adjusting for mode of delivery, intrapartum antibiotic exposure, race and ethnicity, gestational age at birth, and well child visit count before 2 years of age. We did not identify any associations between feeding type in the first six months of life and AOM in the first two years of life (**Figure 4**).

**Figure 3.**
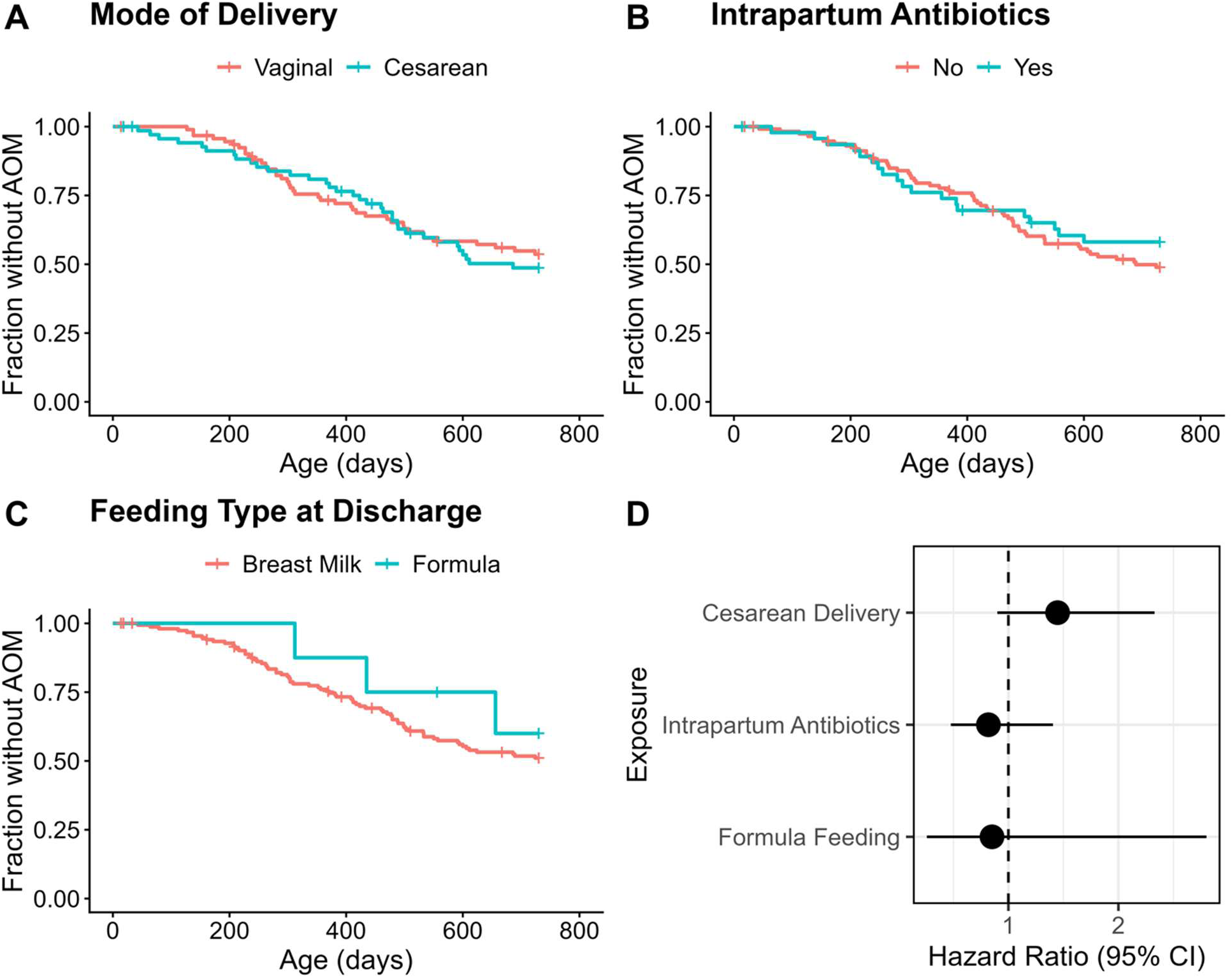
Associations between peripartum exposures and time to first AOM. We constructed Cox proportional hazards models to evaluate associations between time to first AOM episode and (**A**) mode of delivery (vaginal birth n = 93, cesarean section n = 70), (**B**) intrapartum antibiotic exposure (unexposed n = 116, exposed n = 47), and (**C**) feeding type at time of hospital discharge (any breastmilk received: n = 155; exclusively formula fed: n = 8). The survival curves show the proportion of the study population that remains without an episode of AOM across the first 2 years of life. (**D**) The forest plot shows the hazard ratios and 95% confidence intervals for the Cox proportional hazards models of each early life exposure.

**Figure 4.**
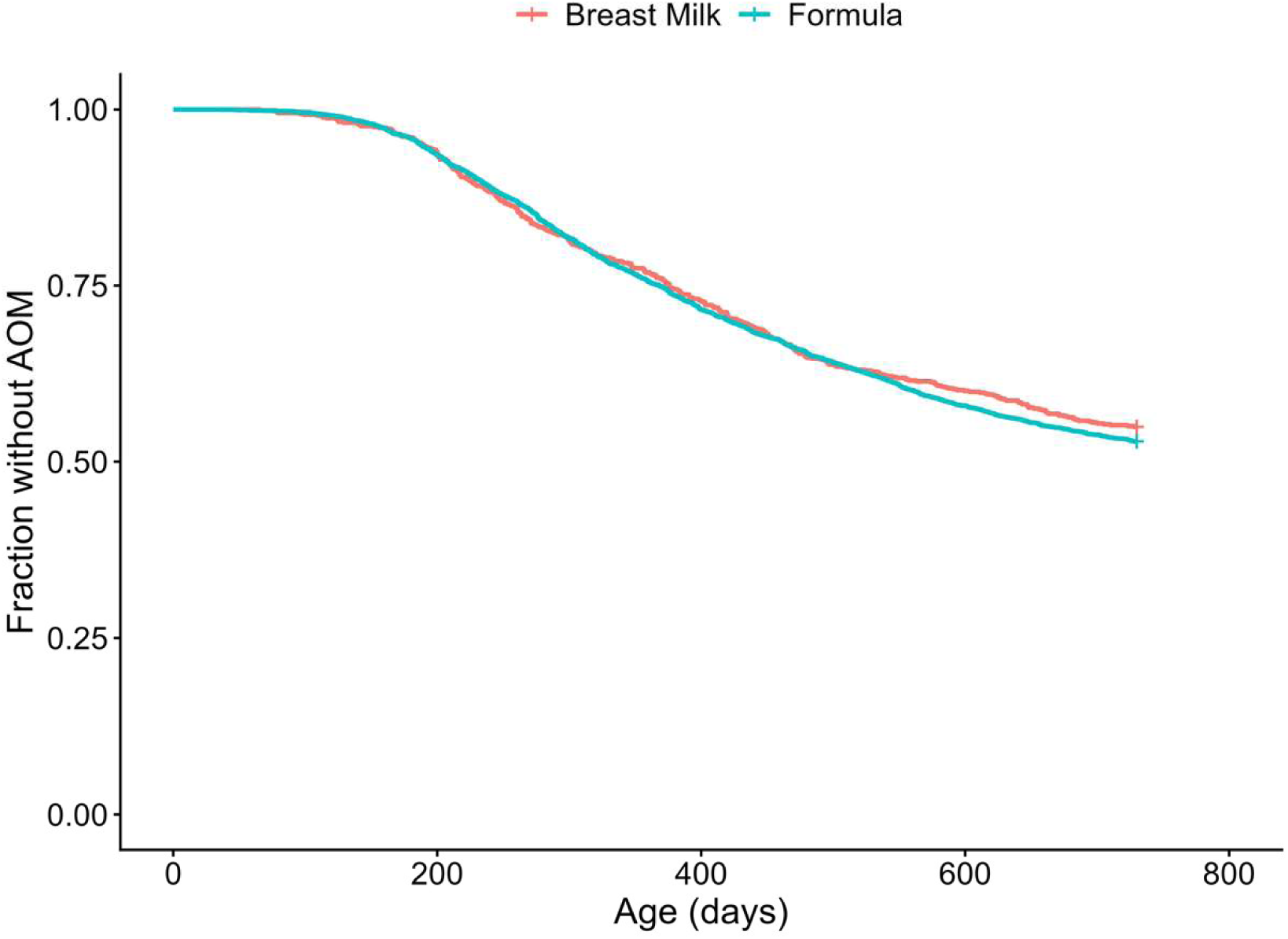
The influence of breastfeeding on time to first AOM episode. We constructed a Cox proportional hazards model to evaluate the association between infant feeding type in the first 6 months of life (any breastmilk received: n = 3,575; exclusively formula fed: n = 803) and time to first AOM episode in infants born at DUHS from January 1, 2021, to June 1, 2023. The survival curves show the proportion of the study population that remains without an episode of AOM across the first 2 years of life.

### Associations between URT microbiome features and time to first AOM episode

To determine if features of the URT microbiome were associated with the time to first AOM episode, we evaluated associations between microbiome richness, Shannon Diversity index, and detection of genera of interest. Cox proportional hazards models that were adjusted for income, NICU admission, and well-child visit attendance. We identified a significant association between the Shannon diversity index and time to first AOM episode, with an increase in Shannon diversity associated with a shorter time to first AOM episode (hazard ratio (HR) (95% CI) 1.40 (1.01 – 1.94), p = 0.041, **Figure 5A**). We did not identify any associations between time to first AOM episode and ASV richness or the presence of genera of interest (**Figure 5B-H**).

**Figure 5.**
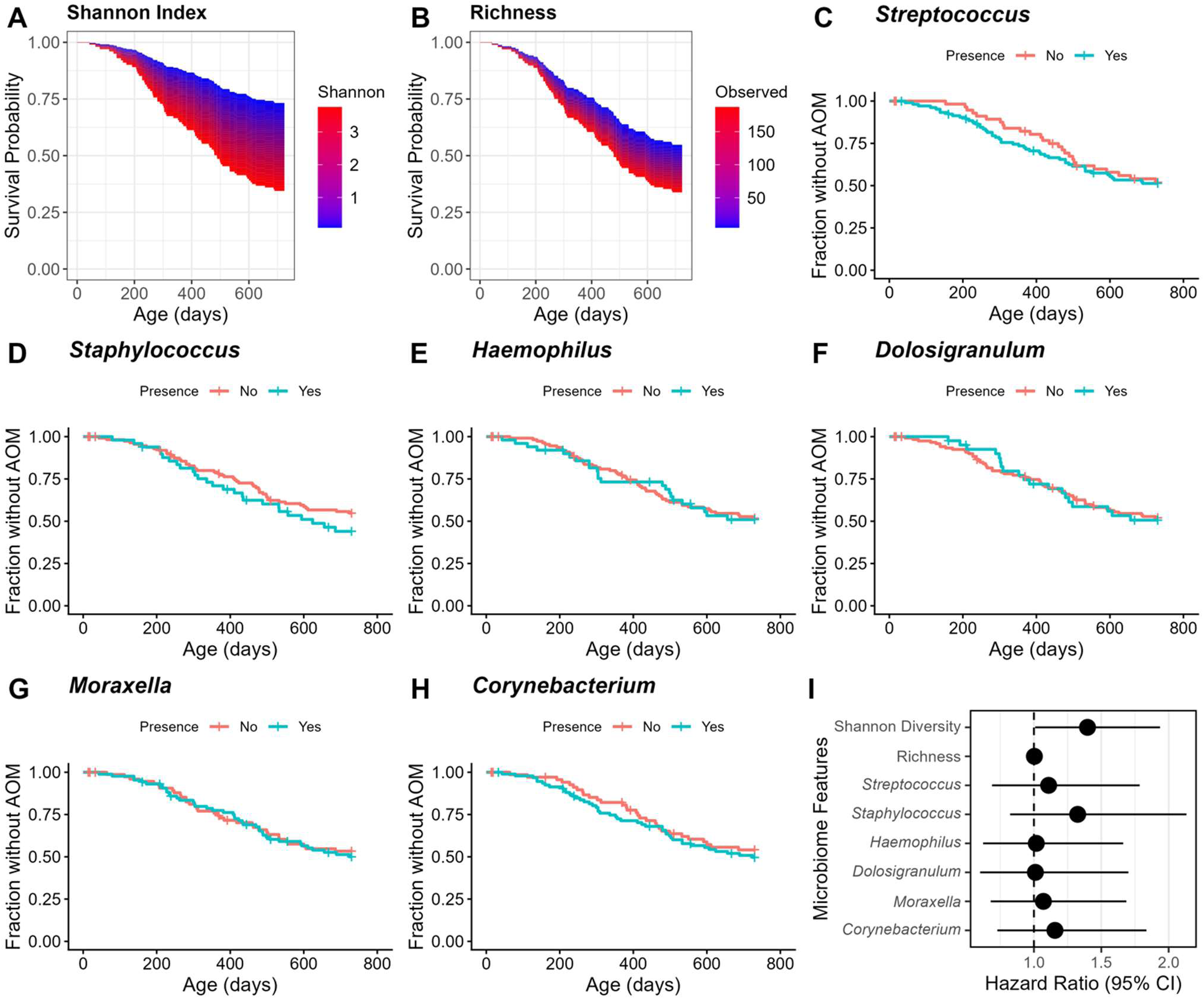
Associations between URT microbiome features, and time to first AOM episode in the first two years of life. We constructed Cox proportional hazards models to determine the associations of key birth microbiome features with the time to first AOM episode within the first two years of life. The survival curves show the proportion of the study population that remains without an episode of AOM across the first 2 years of life for continuous values of (**A**) Shannon diversity index and (**B**) richness. Subsequent survival curves display this proportion across time between groups determined by the presence or absence of genera of interest, including (**C**) *Streptococcus* (present: n = 105 [64%]), (**D**) *Staphylococcus* (present: n = 49 [30%]), (**E**) *Haemophilus* (present: n = 50 [31%]), (**F**) *Dolosigranulum* (Present n = 41 [25%]), (**G**) *Moraxella* (Present n = 86 [53%]), and (**H**) *Corynebacterium* (Present n = 93 [57%]). (**I**) The forest plot shows the hazard ratios and 95% confidence intervals for the Cox proportional hazards models for each of the microbiome characteristics.

### URT microbiome profiles are associated with differences in peripartum exposures and time to first AOM episode

To further characterize differences in URT microbiome composition related to early life exposures, we used unsupervised clustering to classify URT samples into two distinct microbiome profiles (**Figure 6**): a *Streptococcus-*dominant profile (Cluster 1; n = 32) and a diverse profile lacking dominant taxa (Cluster 2; n = 131). We evaluated differences in infant characteristics for each microbiome profile (**Table S5**). The *Streptococcus*-dominant profile was associated with a significantly increased median age at collection compared to the diverse profile (*Streptococcus*-dominant, median (IQR): 32 (24, 49); diverse, median (IQR): 23 (15, 40); Wilcoxon rank sum, p = 0.008) and an increased proportion of infants born via C-section (*Streptococcus*-dominant proportion detected: 59%, diverse proportion detected: 39%; Fisher’s exact test; p = 0.036). We did not identify any associations between cluster and time to first AOM episode (**Figure 6B**).

**Figure 6.**
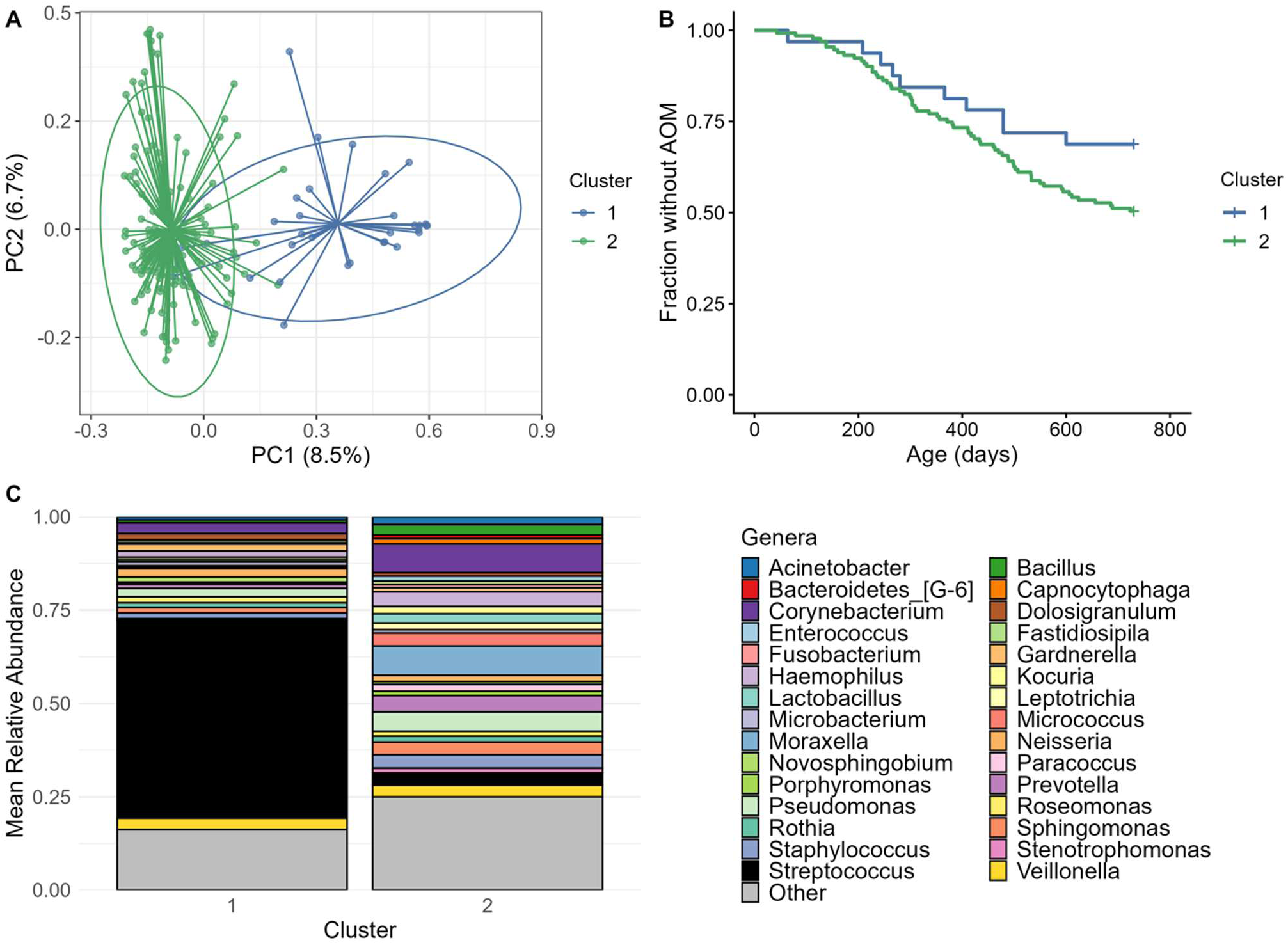
Birth microbiome profiles are not associated with the time to first AOM. We conducted unsupervised clustering of the birth microbiome profiles to evaluate common features between profiles. (**A**) The principal coordinate (PC) plot displays the Euclidean distances and clustering of URT microbiome profiles. (**B**) A Cox proportional hazards model was used to evaluate associations between microbiome profile and time to first AOM episode in the first two years of life. The survival curve shows the proportion of the study population that remains without an episode of AOM across the first 2 years of life for each cluster (Cluster 1 n = 32, Cluster 2 n = 131). (**C**) A stacked bar displays the relative abundance of the top 30 genera identified in the URT microbiome by cluster.

## Discussion

AOM is the most common bacterial infection of childhood. Previous studies have reported increased risk of AOM to early life exposures, including birth via Cesarean section, intrapartum antibiotic exposures, and exclusive formula feeding (8, 10, 14). There is mounting evidence that the URT microbiome is involved in the development of AOM (29, 32, 46). Our study aimed to delineate the influence of these early life exposures on the URT microbiome at birth and to assess for associations with time to first AOM episode. We found that infants who were exclusively formula fed at hospital discharge had increased microbial alpha diversity and increased prevalence of *Haemophilus* spp. We additionally found that an increased alpha diversity at birth was associated with a shortened time to first AOM episode. Finally, we found that *Streptococcus*-dominated microbiome profiles were associated with birth via Cesarean section. These findings suggest that perinatal exposures influence the composition of the URT microbiome at birth and that birth URT microbiome diversity may contribute to AOM susceptibility in infancy.

Breastfeeding in the first several months of life is purported to protect against AOM and respiratory infections in early life, but the specific mechanisms through which breastfeeding mediates protection are still being explored (13, 14, 19). We identified associations between feeding type in early life and microbiome composition, including increased richness and alpha diversity in exclusively formula-fed infants compared to those receiving any amount of breastmilk at the time of hospital discharge. These findings align with those of Rosas-Salazar et al., who found that infants who were exclusively formula-fed had an increased URT microbiome richness and alpha diversity during the first several weeks of life (47). Given that we observed decreased diversity in breastfed children, our findings also join previous studies that suggest breastfeeding in the first weeks of life may delay diversification of the infant gut and URT microbiomes (48, 49). We also observed that formula feeding was associated with increased detection of *Haemophilus* and *Staphylococcus*. Previous studies have shown that early colonization by *H. influenzae* is a risk factor for AOM in early life (30, 46, 50). Similarly, Biesbroek et al. found that formula-fed infants had an increased abundance of *Staphylococcus* at 6 weeks of age and decreased abundances of *Dolosigranulum* and *Corynebacterium* in the URT microbiome, though these differences resolved by 6 months of age (19). Together, these findings suggest that breastfeeding may play an important role in the early seeding and maturation of the infant URT microbiome. Despite observed differences in URT microbiome composition, we did not find an association with feeding type at discharge and time to first AOM episode in the first two years of life within HOPE 1000 infants. Additionally, we did not see an association between breastfeeding in the first 6 months of life and time to first AOM episode in a larger cohort of infants born at DUHS. Our findings contrast previous studies that have found protective associations between breastfeeding and AOM, particularly breastfeeding occurring in the first 6 months of life (13–15). Our results prompt further study into the influence of breastfeeding on mechanisms underlying AOM susceptibility in early life, including the developing immune system and URT microbiome.

Microbial diversity of the URT has previously been associated with protection against AOM and other upper respiratory infections in early life. Xu et al. found that AOM-prone children had a lower URT alpha diversity at 6 months of age compared to children who were AOM-free (29). Additionally, Santee et al. found that increased Shannon diversity was associated with an increased time to first upper respiratory infection (51). In contrast, we found that a higher Shannon diversity at birth was associated with a decreased time to first AOM episode. We are not aware of any studies that have assessed the relationship between URT microbiome diversity at birth and AOM. These contrasting findings suggest that microbiome diversity may not represent a broadly protective quality of the URT microbiome throughout development and/or that birth microbiome may be differentially associated with respiratory health, prompting further analysis of birth microbiome diversity and infection risk.

In addition to evaluating individual features of the URT microbiome, we also evaluated how microbiome composition relates to perinatal exposures. We found that a *Streptococcus*-dominated microbiome profile was associated with older age at time of sample collection and birth via Cesarean section. A study conducted in pre-term infants (gestational ages of 34-36 weeks) found a greater abundance of *Streptococcus* in infants born via cesarean section (mean relative abundance *Streptococcus*: 26.2%) compared to those delivered vaginally (mean relative abundance *Streptococcus*: 3.8%) (52). Similarly, a study conducted by Bosch et al. found that infants born via cesarean section had greater relative abundance of both *Streptococcus salivarius and Streptococcus viridans* compared to infants delivered vaginally (17). Notably, early colonization with the pathobiont *S. pneumoniae* is associated with increased risk for AOM in early life; however, commensal *Streptococci* frequently are associated with protection from infection (29, 53). Because we are not able to resolve taxa to the species level, we cannot determine whether these microbiomes are dominated by pathobiont or commensal *Streptococcus* species and future studies using higher-resolution methods will be needed to provide further insights.

Our study has several strengths and limitations. Samples and data from our prospective birth cohort allowed integration of microbiome specimens with electronic health records data for accurate identification and linkage of medically diagnosed AOM episodes and microbiome features. Due to the timely biospecimen collection at delivery visits, our study provides one of the earliest characterizations of the URT microbiome in infancy. Our study also faced several limitations. We relied on V4 16S rRNA gene amplicon sequencing for characterization of the URT microbiome, which generally lacks species-level resolution. Further analysis with high-resolution sequencing methods, such as shotgun metagenomic or 16S rRNA long-read sequencing, could provide a more comprehensive analysis of putative pathobionts or commensal species within the URT microbiome. We found that formula feeding at time of discharge was uncommon within our study population, with only 5% of infants reported as exclusively formula fed at hospital discharge. While data from our EHR-derived cohort allowed us to evaluate the relationship between infant feeding practices and time to first AOM episode, a larger cohort with additional formula-fed infants and nasopharyngeal swab samples will be needed to evaluate associations between infant feeding and microbiome features. Finally, we relied on EHR documentation of infant feeding practices and AOM episodes, which cannot fully account for interindividual differences in healthcare utilization, diagnoses, or clinical documentation.

Our study provided an in-depth characterization of the birth URT microbiome in a large cohort of infants. This represents an understudied phase in the development of the URT microbiome, an aspect of host physiology that may influence the risk of AOM and other respiratory infections in early childhood. We found that exclusive formula feeding is associated with increased alpha diversity of the birth URT microbiome and greater prevalence of *Haemophilus* and *Staphylococcus* spp. We also found that increased URT microbiome diversity at birth may be associated with decreased time to first AOM episode. These results highlight the role of early life exposures in the development of the URT microbiome and indicate that microbiome-targeted interventions in the perinatal period could potentially be used to improve infant respiratory health.

## Acknowledgments

The authors would like to thank the HOPE 1000 participants. This work was supported by a National Institutes of Health Career Development Award to JHH (K01-AI173398), the Derfner Foundation, and the Department of Pediatrics at Duke University School of Medicine. The content herein is the responsibility of the authors and does not necessarily reflect the views of the funding bodies. The funding bodies had no role in the study design, data collection and analysis, decision to publish, or preparation of the manuscript.

## Data Availability

The sequencing data set used in the completion of this study is available in the Sequence Read Archive (PRJNA1481259). The metadata and code resources associated with this study are available at https://github.com/HurstLab/Hope_Birth_Microbiome_Shortread16S.

